# Towards optimization of oscillatory stimulation during sleep

**DOI:** 10.1101/2021.09.27.461932

**Authors:** Julia Ladenbauer, Liliia Khakimova, Robert Malinowski, Daniela Obst, Eric Tönnies, Daria Antonenko, Klaus Obermayer, Jeff Hanna, Agnes Flöel

## Abstract

**Background:** Oscillatory rhythms during sleep such as slow oscillations (SO) and spindles, and most importantly their coupling, are thought to underlie processes of memory consolidation. External slow oscillatory transcranial direct current stimulation (so-tDCS) with a frequency of 0.75 Hz has been shown to improve this coupling and memory consolidation, however, effects varied quite markedly between individuals, studies, and species. Here, we aimed to determine how precisely the frequency of stimulation has to match the naturally occurring SO frequency in individuals to optimally improve SO-spindle coupling. Moreover, we systematically tested stimulation durations necessary to induce changes.

**Methods:** We addressed these questions by comparing so-tDCS with individualized frequency to standardized frequency of 0.75 Hz in a within-subject design with 28 older participants during napping while systematically varying stimulation train durations between 30 s, 2 min, and 5 min.

**Results:** Stimulation trains as short as 30 s were sufficient to modulate the coupling between SOs and spindle activity. Contrary to our expectations, so-tDCS with standardized frequency indicated stronger aftereffects with regard to SO-spindle coupling compared to individualized frequency. Angle and variance of spindle maxima occurrence during the SO cycle were similarly modulated.

**Conclusion:** In sum, short stimulation trains were sufficient to induce significant changes in sleep physiology allowing for more trains of stimulation, which provides methodological advantages and possibly even larger behavioral effects in future studies. With regard to individualized stimulation frequency, further options of optimization need to be investigated, such as closed-loop stimulation to calibrate stimulation frequency to the SO frequency at time of stimulation onset.

**Significance statement:** Application of slow oscillatory transcranial direct current stimulation during sleep has been shown to enhance specific memory-relevant sleep parameters and memory performance after sleep, albeit with a high degree of variability. Here, we systematically explored two major stimulation parameters possibly accounting for this variability in humans: frequency and duration of stimulation. We found, contrary to our expectations, standardized frequency stimulation with 0.75 Hz being superior to individualized frequency stimulation in enhancing specific sleep parameters. Moreover short stimulation trains of 30 seconds were as effective as 5 min in modulating aftereffects. These are encouraging findings, implying methodological advantages as larger quantity of aftereffects data can be obtained within the same time window, which may also lead to enhanced behavioral stimulation effects.

## Introduction

Sleep is involved in transformation of labile into long-lasting memories. Influential hypotheses postulate that communication between the hippocampus and neocortical networks is pivotal for the consolidation of newly acquired information as consolidation relies on the redistribution of reactivated memories from hippocampus to neocortical long-term storage ^1,2^. This communication may be accomplished by an interplay between neuronal oscillations during nonrapid eye movement (NREM) sleep, namely cortical slow oscillations (SOs), thalamo-cortical spindles and hippocampal ripples ^3,4^. SOs reflect global alterations between neuronal silence (i.e. hyperpolarization; termed down-states) and neuronal excitation (i.e. depolarization; up-states) with a frequency < 1 Hz ^5^, and were shown to have synchronizing effects on cortical as well as subcortical faster oscillatory rhythms such as spindles and hippocampal ripples ^3,6^. The prominent role of SOs orchestrating spindle activity was further underscored, as SO-spindle coupling was not only shown to predict inter-regional connectivity (i.e. communication), but to also initiate hippocampal ripple activity and alter subsequent transfer of information ^3^. These recent findings emphasize the top-down organizing role of SOs.

Application of oscillatory protocols with frequency <1 Hz (slow oscillatory transcranial direct current stimulation, so-tDCS) during NREM sleep provides a non-invasive method of manipulating cortical SO activity ^7–10^, and improves the precisely timed coupling between endogenous SOs and spindles, as shown in patients with mild cognitive impairment ^8^. Critically, in parallel, so-tDCS improves memory retention performance at group level ^7–11^, with coupling between SO and spindles being most informative for memory retention performance ^8^. Therefore, so-tDCS during sleep not only represents a non-invasive, well-tolerated method to test causal effects of external SO induction on endogenous oscillatory rhythms and its relation to memory, but may also provide a promising treatment method in aging individuals with memory decline in the future. However, response to stimulation on electrophysiological and behavioral levels varied quite markedly between individuals ^8–10^, studies ^9,10,12,13^ and species ^14,15^. Some studies did not find an overall effect ^13,16^. Such inconsistent effects may be due to – among other reasons – inter-individual differences in brain traits and states, which might not respond equally well to a standardized so-tDCS frequency of 0.75 Hz. Previous evidence from transcranial alternating current stimulation (tACS) studies suggests that oscillatory stimulation is most effective if stimulation frequency is matching the network’s preferred frequency (eigenfrequency), as in that case intrinsic rhythmic activity is assumed to adapt more easily to the external stimulation ^17,18^. However, it is unclear how eigenfrequency so-tDCS will affect human sleep physiology, in particular SO-spindle coupling as the memory-relevant measure.

Furthermore, so-tDCS train duration might affect the magnitude of aftereffects in terms of SO-spindle coupling, but so far, effects of stimulation duration have not been systematically explored.

To address these open questions, we tested potential differences between so-tDCS with eigenfrequency and standardized frequency of 0.75 Hz on SO-spindle coupling during napping; and systematically explored differences between diverging lengths of stimulation trains (30 s, 2 min and 5 min) in 28 older adults using a cross-over design. We hypothesized that eigenfrequency stimulation would induce strongest stimulation effects. Moreover, we hypothesized that effects would be most pronounced for longest stimulation duration (5-min trains).

## Materials and methods

### Participants

Participants were recruited from the Department of Neurology, Universitätsmedizin Greifswald, Germany (ClinicalTrials.gov Identifier: NCT04714879). Fifty-four healthy older participants between 50 and 79 years (M = 66.09, SD = 8.09) were included in the study after an initial structured telephone interview to clarify exclusion criteria. Eligible participants then underwent baseline assessment including medical exam, cranial MRI, neuropsychological testing, multiple questionnaires on sleep quality and habits (see Supplementary Information for more details and Table 1 for baseline characteristics). Twenty-eight participants were included in the final analyses, after exclusion due to insufficient sleep in multiple nap sessions (n=14), too few SO events (n=3) or other reasons (e.g. cognitive impairment, sleep apnea, eigenfrequency at 0.75 Hz, disqualifying medication; n=9), for details see Supplementary Information.

**Table 1.** Baseline characteristics.

|  |  |
| --- | --- |
| n (female/male) | 28 (15/13) |
| Age (y) | 65.3 ± 8.5 (50-77) |
| Education (y) | 15.3 ± 3.1 (12-24) |
| AVLT, learning ability | 60.5 ± 6.5 (46-72) |
| AVLT, delayed recall | 13.1 ± 1.9 (8-15) |
| Rey–Osterrieth Complex Figure (copy) | 35.1 ± 1.4 (31-36) |
| Rey–Osterrieth Complex Figure (delayed recall) | 22.9 ± 5.8 (13-33) |
| Stroop color-word test (delay for incongruent vs neutral condition) | 53.9 ± 20.1 (-3 - 91) |
| Verbal fluency, phonematic (no. of words) | 13.9 ± 4.3 (7-24) |
| Verbal fluency, categories (no. of words) | 24.9 ± 5.4 (15-37) |
| Digit span, forward | 8.3 ± 1.9 (6-13) |
| Digit span, backward | 6.7 ± 1.7 (4-11) |
| Letter digit substitution | 29.4 ± 6.0 (16-43) |
| MWT | 32.5 ± 2.3 (28-36) |
| Beck's Depression Index | 3.3 ± 3.5 (0-13) |

Data are given as mean ± SD and range (minimum to maximum). AVLT, Auditory Verbal Learning Test; MWT, Multiple-choice vocabulary intelligence test.

The study was approved by the local ethics committee (Universitätsmedizin Greifswald) and conducted in accordance with the Declaration of Helsinki. Participants gave written informed consent prior to participation.

### Procedure

After an adaptation nap in the sleep laboratory, participants first underwent one nap with EEG measurement to determine eigenfrequency. Participants were then tested in a within-subject design to compare standardized so-tCS with a frequency of 0.75Hz to eigenfrequency so-tCS, while varying three stimulation train durations resulting in seven conditions including sham (see Fig. 1A for Experimental design). Conditions were conducted in counterbalanced order with randomized order assignment and were separated by an interval of at least one week. Participants were blinded for stimulation condition during the complete study. All nap sessions took place at the sleep laboratory of the Department of Neurology, Universitätsmedizin Greifswald, Germany. Upon arriving between 11.30 a.m. and 2 p.m. depending on their daily routine, participants were prepared for EEG-recordings. EEG was recorded using the BrainAmp amplifier system (Brain Products, Munich, Germany) and an actiCAP equipped with (at least) 26 active recording electrodes. After a standardized small meal and preparation for stimulation, participants were asked to attempt to sleep during a period of 90 minutes. Oscillatory anodal current stimulation was applied during stable NREM sleep stage 2 by two battery-driven stimulators (3–5 trains; DC-Stimulator; NeuroConn, Ilmenau, Germany) to bilateral frontal electrodes (8mm in diameter; max. current density of 0.522 mA/cm^2^ in each hemisphere, reference electrodes placed at each mastoid) with a stimulation protocol similar to the one described previously by Ladenbauer et al. ^8^. The stimulation frequency was 0.75Hz in the standardized stimulation condition and between 0.5 and 1Hz in the eigenfrequency stimulation condition, which was determined using the SO power peak by means of the Gaussian fit approach. For more details on sleep monitoring, sleep scoring, stimulation and assessment of eigenfrequency see Supplementary Information.

**Figure 1.**
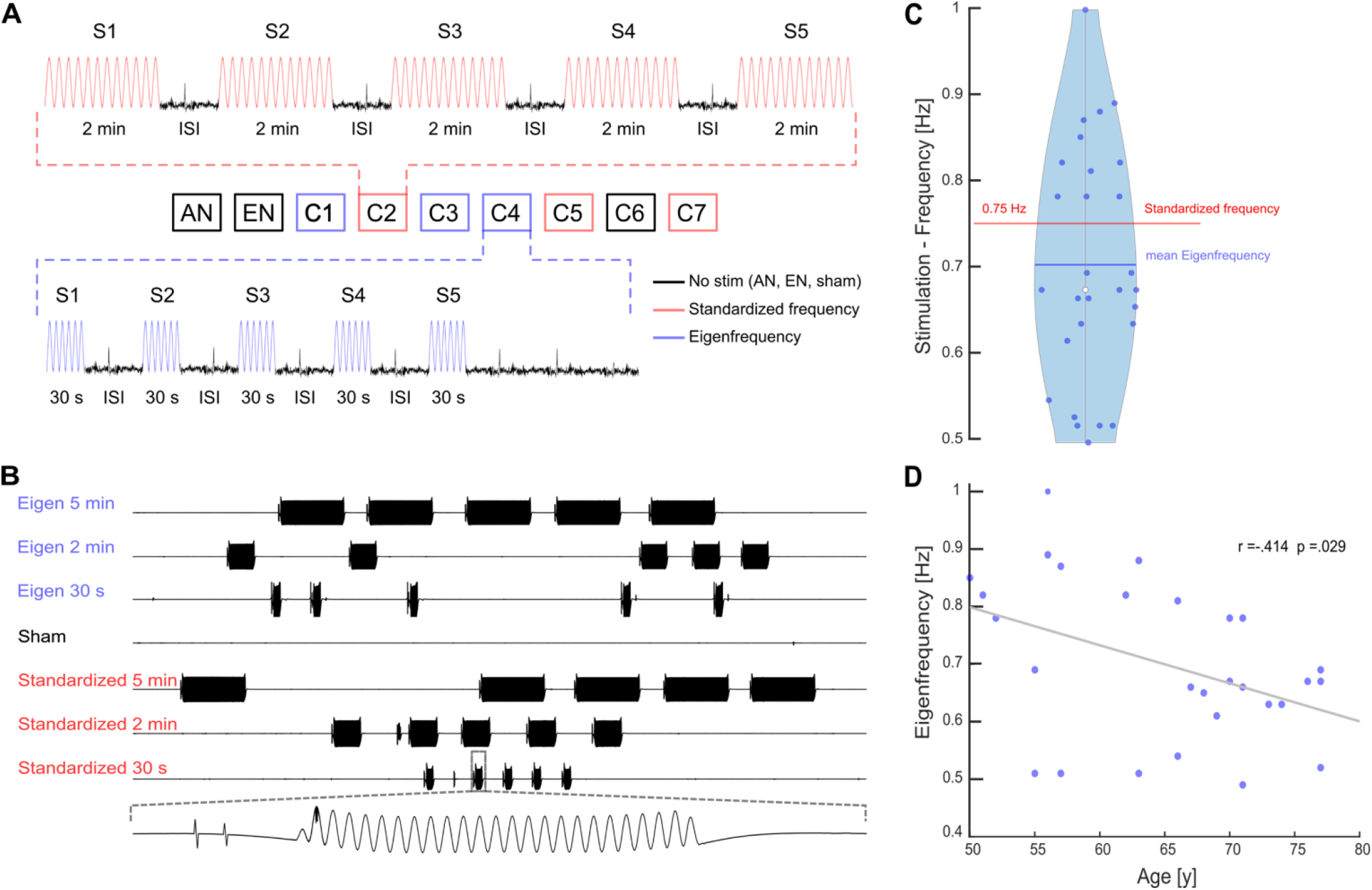
Experimental design and eigenfrequency. (A) Graphic illustration of the experimental design. Each participant underwent 9 nap sessions. First, participants attended a habituation nap (HN) in which they were familiarized with the experimental setup and procedures (following baseline assessment on the same day). The second ‘eigenfrequency nap’ (EN) session served to determine individual SO peak frequency. The following 7 experimental nap sessions consisted of 6 so-tDCS conditions, in which eigen-(blue) and standardized (red) frequency was paired with each stimulations duration (30 s, 2 min, 5 min), and one sham condition (black). (B) Example nap sessions for each condition. Black clusters represent individual stimulation trains. Number of stimulation trains (3–5) and duration of interstimulus intervals between stimulation trains depended on individual partcipant’s sleep, as each train was only initiated during N2 or SWS only. (C) Overview of stimulation frequencies. Blue dots indicate individual eigenfrequencies, a blue line represents the mean and a white dot the median. Standardized stimulation is visualized as a red line. (D) Scatterplot showing the association between age and eigenfrequency.

### SO detection

After preprocessing (see Supplemantary materials for detailed information), analysis was constrained to the average of six fronto-central channels (Fz, FC1, FC2, Cz, CP1, and CP2), based on findings in past research ^8,9,19^. Detection of SOs also mostly followed practices established in the literature (e.g. ^6,8^). In brief, the averaged fronto-central signal was filtered with a finite impulse response bandpass of 0.16-1.25Hz. Then all negative-to-positive zero-crossings in the signal were identified. The time between two successive negative zero-crossings was measured, as well as the amplitude difference between the most negative point (down-state trough) after the first positive-to-negative zero-crossing, and the most positive point (up-state peak) after the next negative-to-positive crossing. If the time between two successive negative-crossings was between 0.8-2s (0.5-1.25 Hz) and amplitude difference within the 65^th^ percentile of all amplitude differences, then the event was marked as a SO.

### Time-frequency representation analysis

Because we were exploring the effects of a wide range of stimulation parameters, we opted to use several complementary approaches to attain a fuller picture of the effects of different types of stimulation on SO-spindle coupling. We started with a Time-Frequency Representation (TFR) analysis ^20^, as it can be used in concert with linear mixed effects (LME) models, thus allowing a rigorous statistical analysis that provides an overview of what stimulation parameters most affect brain measures, and at what times/frequencies, while also being able to factor out nuisance variables.

TFR were calculated within the spindle band (10-20 Hz) for each SO event (5-sec epoch) using a morlet wavelet transform with 5 cycles, after data were first down-sampled to 50 Hz. The first and last 150 ms of each epoch were cut off to remove edge effects. Baseline adjustment followed the z-score method and correction for multiple comparisons was conducted with the permutation clustering approach (please see Supplemental information for more details). Clustering was then applied to the main LME test statistics (see Supplemental information).

### Linear mixed effects models

Event-related neurophysiological data have traditionally been analyzed by averaging the events by condition and participant. With enough events, this achieves the effect of canceling out noise, and produces a single value per condition and subject, which can then be submitted to commonly-used parametric tests. This approach is simple, and computationally tractable, but also has disadvantages (see ^21^ and ^22^ for discussion), primarily the destruction of all information pertaining to within-subject variance. To avoid these issues and to produce more robust statistical inferences, we applied LME models. Rather than averaging across trials per participant, each individual trial was entered as a data point.

The fixed effects for the LME models included so-tDCS stimulation type (sham, eigen-, or standardized frequency), and stimulation duration (30 s, 2 min, 5 min). The random effects included participant, and the interaction of stimulation synchronicity with stimulation (see Supplemental information) and duration. In the case of TFR analysis, a separate model was fit for each point in 2D time-frequency space.

### Event-Related Phase Amplitude Coupling

Given that in TFR analysis phase is only indirectly measured and alignment of time and SO phase somewhat imprecise, we also analyzed the data with Event-Related Phase Amplitude Coupling (ERPAC) ^23^. Here, the correlation of phase and power is directly computed at each time/frequency point, across trials, which provides a more fine-grained picture of the phase-power relationship. It is however not straightforward to compare several, interacting factors; so here it was useful to be able to submit a simplified analysis to ERPAC constrained on the basis of the TFR analysis.

ERPAC analysis was carried out with the Python package Tensorpac 0.6.5 ^24^. Briefly, for each time-frequency point a regression is performed across trials with power in the power band of interest as the regressand, and the sine and cosine of the phase in the phase band of interest as regressors. This produces a rho and corresponding p-value for each point in time-frequency space, allowing inferences on which times and frequencies there is a reliable relationship between phase and power. Separate ERPAC analyses were performed for sham, eigen-, and standardized frequency conditions. All trials from all participants from a given condition were used; within-subject variance was not estimated in the ERPAC analysis. Differences between the standardized stimulation and sham, and between eigenfrequency stimulation and sham were compared according to the procedure used in ^23^; for details see Supplementary information.

### Phase of spindle maxima

The two above mentioned methods are sensitive to the presence of spindles nested in SOs, but cannot say anything about the spindle dynamics themselves. To explore these, we further measured the phase of the SOs where spindle band power reached its maximum, similarly to previously established procedures ^8,25^. The fronto-central channel signal in the slow oscillations was transformed into instantaneous phase in the 0.5-1.25Hz band with a wavelet transformation. The same signal was then transformed into power, also with a wavelet transformation, in two bands: 12-15 Hz and 15-18 Hz. In each power band, for each SO, the time point where power was maximal was identified, and the phase at this time point was selected as the SO phase of maximal power. Differences in distributions of phases across the sham, eigen-, and standardized frequency conditions were summarized with their resultant vectors (see Supplementary Information for more details), and compared across conditions with the two sample Kuiper test, which is the circular analogue of the Kolmogorov-Smirnoff test. Here, as with ERPAC, a complex, factorial statistical analysis is not possible, so we simplified the analysis with constraints provided from the results of the TFR analysis. Signal decomposition was carried out with the Python package Tensorpac 0.6.5 ^24^.

## Results

With regard to the detected eigenfrequency in our sample of healthy older adults, we found 17 participants exhibiting an SO peak below 0.7 Hz, and tested for an association with age. Figure 1B/C shows the distribution of eigenfrequencies and association with age. Eigenfrequency was negatively correlated with age (r= −0.41, p= 0.029).

Table 2 depicts the number of stimulation trains that participants received in each condition. Slightly more stimulation trains were applied for standardized as compared to eigenfrequency stimulation for all three durations, however, these differences were not significant (all p’s > 0.1; please see Table 2). Only during the sham session were significantly more sham blocks possible, when 30 s or 2 min sham trains were scored applying same criteria as for so-tDCS conditions (Z = −2.428, p= 0.015 and Z= −2,070, p= 0.038, respectively; Wilcoxon signed-rank tests).

**Table 2.** So-tDCS/sham train count.

| **30-s condition (number of participants)** |  |  |  |
| --- | --- | --- | --- |
| Nr. of stimulation trains | eigenfrequency | standardized frequency | sham stimulation |
| 5 | 21 | 26 | 28 |
| 4 | 3 | 1 | 0 |
| 3 | 4 | 1 | 0 |
| **2-min condition (number of participants)** |  |  |  |
| Nr. of stimulation trains | eigenfrequency | standardized frequency | sham stimulation |
| 5 | 23 | 24 | 28 |
| 4 | 3 | 3 | 0 |
| 3 | 2 | 1 | 0 |
| **5-min condition (number of participants)** |  |  |  |
| Nr. of stimulation trains | eigenfrequency | standardized frequency | sham stimulation |
| 5 | 21 | 23 | 22 |
| 4 | 4 | 4 | 5 |
| 3 | 3 | 0 | 1 |

Similarly, with regard to macro sleep structure, no differences were observed for standardized versus eigenfrequency stimulation conditions (all p> 0.27; Welch ANOVAs; please see Supplementary Information Table 2-1 for sleep structure in each condition). A significant difference was only observed for the wake time during the sham session in comparison to the 2-min standardized frequency condition (Welch-ANOVA (*F* (2, 46.501)= 4.710, p=0.014, p= 0.023; Bonferroni-corrected), which may be due to one outlier of 42 min wake time during the sham session.

Detection algorithms identified 6964 slow oscillations during the 1-min stimulation free intervals over all conditions. These oscillations and their accompanying spindle band power were analyzed here with several, complementary forms of phase amplitude coupling analyses.

### Time Frequency Representation

In order to obtain an overview of phase power relationships with an exhaustive statistical analysis, we first compared time-frequency representations in the spindle band for SOs under varying stimulation conditions. Differences between conditions were inferred using mass-univariate LME models. The 30s sham condition was chosen as the baseline state, or intercept. The model parameters/predictions indicate the estimated changes that stimulation type and duration had on the baseline state. After correcting for multiple comparisons, there was a general effect of standardized frequency stimulation in the ~15-18 Hz frequency band, 90-180 ms after the down-state trough, such that power increased during the late rising phase of the SO cycle. Notably, the interactions of 2 min and 5 min with standardized stimulation frequency were not significant, indicating this general standardized frequency effect was not reliably modulated by stimulation duration. No significant cluster was found for eigenfrequency (see Fig. S1).

Model predictions are depicted in Figure 2 and 3A – these are analogous to condition averages, and can be interpreted the same way. Parameter estimates and their corresponding significance are depicted in Supplementary Figure 1.

**Figure 2:**
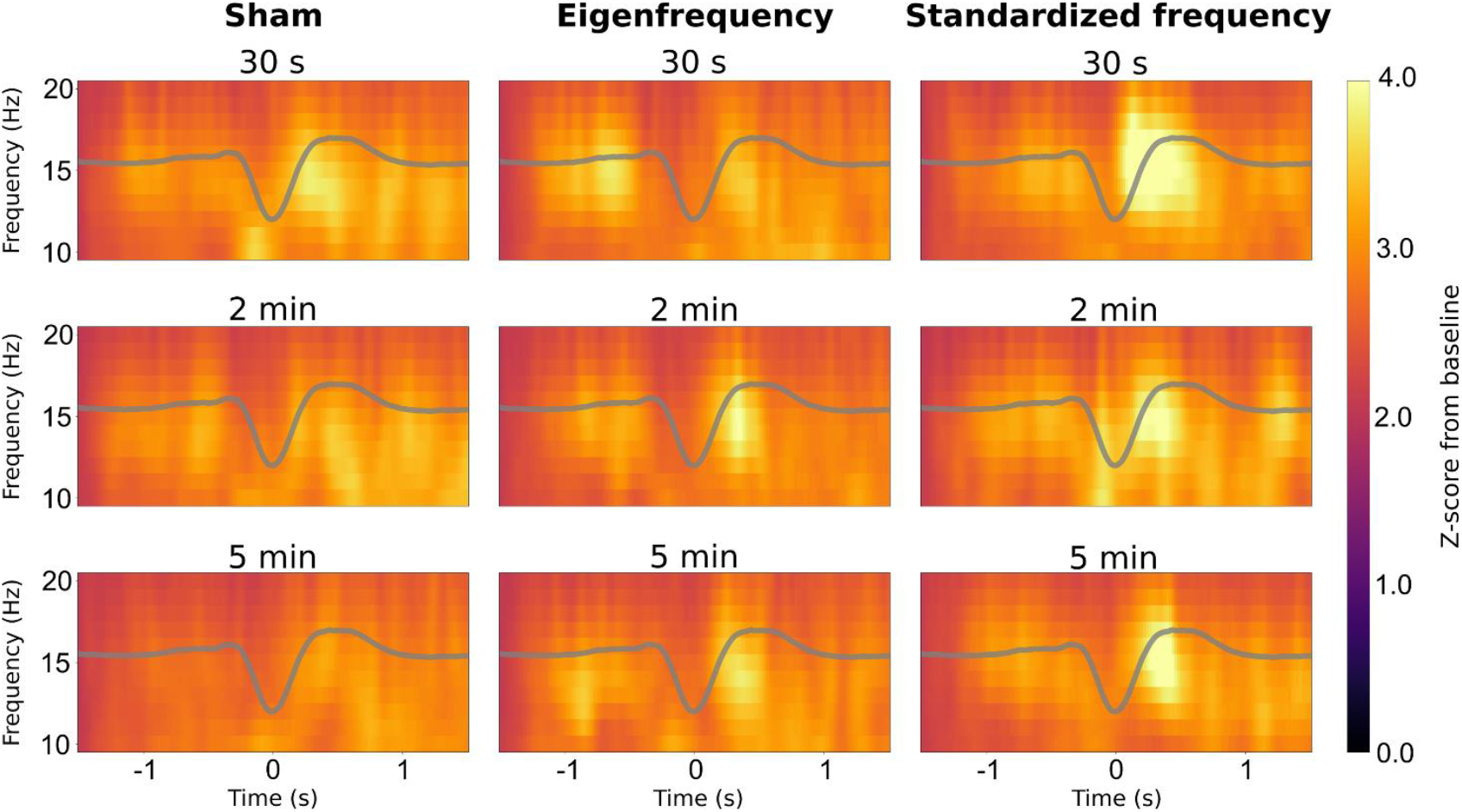
LME model predictions for spindle band power during SOs for each condition. (analogous to condition averages). Averaged slow oscillations are superimposed in gray.

**Figure 3:**
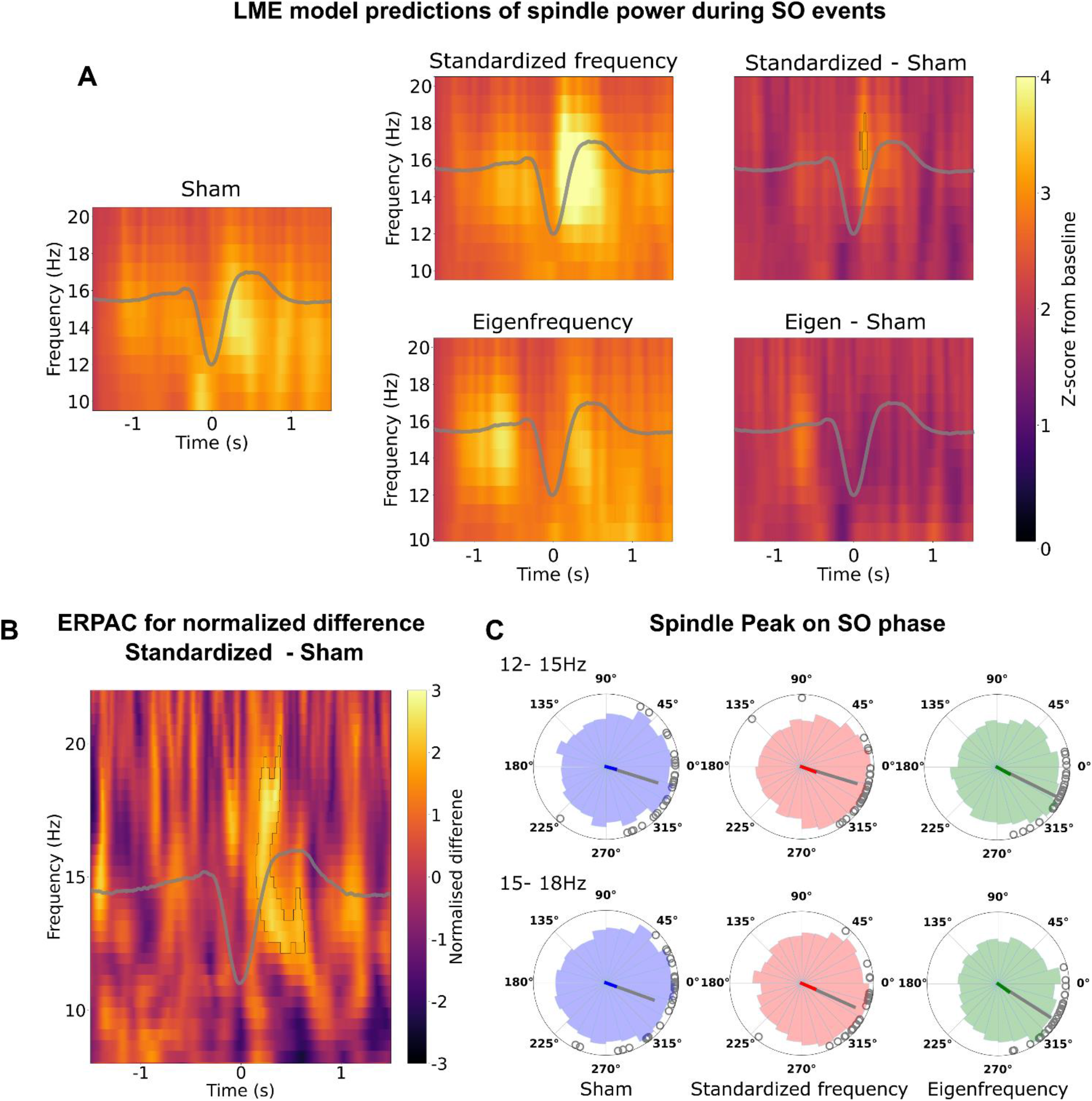
Relationship between SO phase and spindle band power as assessed by several complementary methods. A: LME model predictions for sham (left), eigen-(bottom) and standardized frequency (top) stimulation locked to SO troughs, and the subtractions of sham from the latter two. Statistically significant deviations from sham are indicated with black outlines. Averaged SOs are superimposed in gray. B: Normalized difference between standardized frequency ERPAC and sham ERPAC locked to the SO through. Areas of significant difference, corrected for multiple comparisons at p> 0.05, are indicated with black lines. Averaged slow oscillations are superimposed in gray. C: Polar histograms of SO phase of maximum spindle band power in 12-15 Hz and 15-18 Hz frequency range for sham (left, n= 2600), eigenfrequency (center, n= 2020) and standardized frequency (top, n= 2344) stimulation condition. Colored lines are the resultant vector for all phases; line direction indicates circular mean, and line length is inversely related to circular variance. Gray dots indicate the circular average for individual participants, and the gray line is their corresponding resultant vector.

### Event-related Phase Amplitude Coupling (ERPAC)

Next, we more directly assessed the correlation of phase and power with ERPAC ^23^. Results are depicted in Figure 3B. These show a reliable increase in the relationship between phase and power for standardized stimulation frequency over sham, starting during the ascending up-state (~180-380ms) at 15-18 Hz, and extending throughout the duration of the up-state (~250-650ms) at 12-15 Hz. As a factorial analysis is not possible with ERPAC, and since the TFR did not show any interaction of stimulation with duration, we collapsed the duration conditions and only tested stimulation type.

### Phase of spindle band maxima

The results from TFR and ERPAC analyses seem to converge on the 12-15 Hz and 15-18 Hz bands as loci of spindle band power modulation for the slow oscillations. In order to more directly examine spindle dynamics in relation to *oscillation phase*, as well as assess spindle variance across trials/participants, we compared the phase of slow oscillations at spindle maxima across different stimulation conditions in these two spindle bands (see Fig. 3C). Consistent with past research on older participants, spindles tended to concentrate somewhat earlier than the up-state ^8,25,26^; in contrast, spindles in younger participants have been reported to concentrate directly on the up-state ^25,26^, although not all studies could confirm these findings ^27,28^. Comparison of the phase distributions in the 12-15 Hz band with a two sample Kuiper test found differences between sham and both stimulation conditions (to eigenfrequency: D= 0.063, p= 0.004 and standardized frequency: D= 0.056, p= 0.013), but not between eigen- and standardized frequency stimulation (D= 0.045, p= 0.196). In the 15-18 Hz band, only distributions between standardized and sham condition differed (D= 0.078, p< 0.001), while no differences were evident between other conditions (all p> 0.05).

In addition to overall phase distributions, the distribution of participant mean phases (gray dots in Fig. 3C) were analyzed in the same way. In the 12-15 Hz spindle band no differences between conditions were observed (all p> 0.1), however both eigen- and standardized frequency distributions differed to sham distribution in the 15-18 Hz band (D= 0.571, p= 0.001 and D= 0.536, p= 0.004, respectively), with no difference between the two (D= 0.357, p= 0.234). The length and direction of the resultant vectors of phase (Fig. 3C and Table 3) indicate the general trend that both stimulation conditions, eigen- or standardized frequency, decrease the phase variance in comparison to sham, while pushing the mean phase value in the counter-clockwise direction. Descriptively, standardized frequency stimulation has a slightly stronger effect than eigenfrequency.

**Table 3:** Phases of spindle maxima: resultant vectors.

|  | Resultant vector direction<br>(radians/degrees) | Resultant vector length |
| --- | --- | --- |
| **Phases at power band 12-15 Hz** |  |  |
| Sham | -0.255/345 | 0.19 |
| Eigenfrequency | -0.36/339 | 0.25 |
| Standardized | -0.51/331 | 0.22 |
| **Phases at power band 15-18 Hz** |  |  |
| Sham | -0.35/340 | 0.17 |
| Eigenfrequency | -0.399/337 | 0.22 |
| Standardized | -0.627/324 | 0.23 |
| **Participant mean phases at power band 12-15 Hz** |  |  |
| Sham | -0.304/343 | 0.8 |
| Eigenfrequency | -0.298/343 | 0.83 |
| Standardized | -0.461/334 | 0.93 |
| **Participant mean phases at power band 15-18 Hz** |  |  |
| Sham | -0.345/340 | 0.77 |
| Eigenfrequency | -0.423/336 | 0.87 |
| Standardized | -0.565/327 | 0.93 |

## Discussion

In the current study, we showed that applying a standardized frequency of 0.75 Hz stimulation during NREM sleep can reliably increase the coupling of sleep spindles with SOs, while no effect was observed for eigenfrequency stimulation, as evidenced by an analysis of TFR. Further, this effect seemed to hold with a large variety of stimulation durations, including trains as short as 30 s. An ERPAC analysis, collapsed across durations, found a broader area of spindle enhancement than the TFR analysis for standardized frequency, consisting of a spindle enhancement in the classical fast spindle frequency range (12-15 Hz) during the SO up-phase and a short preceding spindle burst in a higher spindle frequency range (15-18 Hz).

Both frequency conditions reduced the variance of spindle timing within SO phase, while surprisingly pushing their angle slightly towards the late rising phase of the SO rather than to synchrony with SO peak.

### 30 seconds of so-tDCS were sufficient to enhance endogenous SO-spindle coupling

Following the first so-tDCS study during sleep in humans ^7^, the majority of previous studies employed 5-min trains of stimulation to affect SO activity and memory performance ^8–12,29–31^. Animal studies, in contrast, typically used protocols with shorter train durations such as 30 s ^15,32^ or 1 min ^14^ to accommodate for the shorter NREM sleep epochs in rodents, and found online entrainment effects during stimulation, but no significant effects that persisted after stimulation offset. It was therefore suggested that diverging results with respect to these aftereffects are due to differences in stimulation duration, as longer stimulation trains may be necessary to elicit synaptic plasticity changes that persist after stimulation ^33,34^. Here, we systematically investigated shorter compared to longer stimulation train durations and demonstrated for the first time that so-tDCS trains as short as 30s were sufficient to induce lasting effects in humans as evidenced by increased spindle activity during the rising phase of the SO after stimulation offset. In conclusion, 30 s of so-tDCS trains are long enough to cause plastic changes, and aftereffects are even comparable in magnitude with significantly longer stimulation durations.

This finding can be incorporated into future studies, as this offers, on the one hand, methodological advantages by increasing the quantity of after-effects data, and on the other hand possibly also enhances neurophysiological and behavioral stimulation effects in humans.

### Standardized but not eigenfrequency reliably enhanced endogenous SO-spindle coupling

For optimization of oscillatory stimulation protocols, the use eigenfrequency has been repeatedly suggested ^17,35,36^. We therefore tested whether so-tDCS with eigenfrequency rather than standardized 0.75Hz frequency would lead to improved SO-spindle coupling following stimulation.

Contrary to our hypothesis, we found that eigenfrequency so-tDCS did not have reliable effects on triggering and grouping spindles, apart from small changes to phase variance and angle. Standardized frequency of 0.75 Hz, on the other hand, amplified SO-spindle coupling, replicating so-tDCS aftereffects previously reported for 5-min stimulation trains in patients with patients with mild cognitive impairment (MCI)^8^. This finding is unexpected in light of previous literature on stimulation effects following tACS adjusted to eigenfrequency in the targeted (mostly alpha) frequency range ^36–39^. However, not all previous studies have found individualized tACS frequeny to be superior ^40^, or indicated that slight deviations from individual target frequency are favorable ^41^. Vossen and colleagues ^41^ reported a larger effect following stimulation with frequency slightly below the eigenfrequency, and proposed spike-timing-dependent plasticity as one possible explanation: synaptic strengthening within an involved oscillatory neural circuit of a specific frequency band may be achieved when pre-synaptic events at endogenous frequency precede the post-synaptic events induced by tACS. As in our standardized frequency stimulation condition more participants were stimulated with higher SO-frequency in comparison to their eigenfrequency, this explanation seems rather unlikely.

An alternative explanation for these findings may be the specific frequency range 0.7-0.8 Hz that may have greater driving and grouping effects on thalamo-cortical spindles than lower and higher SO sub-frequencies. Young participants ^19,42,43^ as well as cats ^44^ were shown to exhibit pronounced peaks in this narrow frequency range during NREM sleep, possibly indicating an optimal frequency in functional terms. A larger proportion of our older participants, in contrast, indicated an individual SO frequency peak below 0.7 Hz, which was moreover negatively association with age. A descriptively lower SO peak frequency also occurred in a previous study comparing healthy older and young adults (^25^, see Figure 1B EEG power spectra). The authors demonstrated that frontal SOs drive spindle activity only in young adults, and spindles peaked earlier in the rising phase of the SO in older relative to young participants ^25^. This may possibly be explained by an age-related slowdown in SO eigenfrequency. To substantiate this speculative idea, further studies are needed to investigate sub-frequencies of the SO range with respect to their impact on driving spindle activity.

Some methodological aspects with regard to determining eigenfrequency should also be taken into account when interpreting these results. Eigenfrequency was assessed during a previous nap session, but may have been not optimal for the specific nap that was stimulated. A mismatch between stimulation frequency and endogenous SO frequency at time of stimulation may have emerged, that may have hampered so-tDCS online (entrainment) effects and thus so-tDCS aftereffects. A mismatch can even occur despite calibrating stimulation at participants’ individual frequency just prior to the experiment ^36,39,41^. Thus online adjustment of stimulation frequency in a closed-loop approach may be better suited to resolve the question of optimized so-tDCS frequency. Therefore, in the future, the impact of different stimulation frequencies must be explored in a closed-loop approach, possibly aided by neurocomputational models, which address both the generation of SOs in cortical areas ^45,46^ and the interaction between SOs and spindles through the thalamo-cortical loop ^47,48^.

### Limitations

A few limitations of the study should be noted: First, several methodological aspects with regard to determination of an individual’s eigenfrequency may have impacted the current results. Apart from the fact that SO eigenfrequency was assessed during a previous nap session, assessment of individual SO frequency in terms of spectral or event-based approaches may also play a role, as Fourier spectra are not able to discriminate between periodic phenomena (pace) and duration/shape of the two components of SOs ^5^. We used the spectral approach to assess eigenfrequency in accordance with previous literature on individualized alpha frequency ^36,38,39^, but mean frequency of all detected SOs in each participants possibly reflects SO eigenfrequency more accurately.

Second, a technical problem with our stimulation device caused divergence in phase between both hemispheres with increasing stimulation train duration in some of our participants, which needs to be taken into account when interpreting the results. However, stimulation synchronicity was modeled in the LME model as a random effect to statistically exclude the potential variance caused by non-synchronous stimulation.

### Conclusion

In sum, our results indicate no benefit of individualized stimulation frequency relative to standardized 0.75 Hz so-tDCS for the improvement of memory-relevant sleep oscillations in healthy older adults. These results replicated previously reported significant modulation of sleep physiology by 0.75 Hz stimulation in MCI patients and healthy older adults. Our findings further indicate that a stimulation train as short as 30 s is sufficient to increase SO-spindle coupling following so-tDCS with a frequency of 0.75 Hz. From a practical point of view, these findings render stimulation over repeated naps or nights in a non-experimental setting more feasible. Moreover, they open up the possibility to apply several trains of stimulation within one sleep session, possibly inducing larger aftereffects in terms of neurophysiology and memory consolidation. In the future, we will further address individualized stimulation and optimization of so-tDCS during sleep by using online adjustment of stimulation frequency in a closed-loop approach and in interaction with computational modelling methods to systematically test the range of so-tDCS frequencies.

## Authorship statement

JL, KO, JH and AF conceived and designed the study. LK, DO and ET collected data for the study. JH, JL, LK and RM analyzed and interpreted the data. JL, LK and JH wrote the first draft of the manuscript. JL, DA, JH and AF edited and revised the manuscript.

