## Supplementary Materials for "Towards optimization of oscillatory stimulation during sleep"

### Supplementary Methods

#### Participants

We tested older adults as the target population of strategies aiming at improving cognitive health in aging and for establishing a baseline group for future studies on patients with neurodegenerative diseases. The exclusion criteria comprised history of severe untreated medical, neurological, and psychiatric diseases, sleep disorders, alcohol or substance abuse, intake of medication acting primarily on the central nervous system (e.g., antipsychotics, antidepressants, benzodiazepines, or any type of over-the-counter sleep-inducing drugs), cognitive impairment or subjective cognitive decline. During baseline visits, eligible participants underwent a medical and neuropsychological screening including magnetic resonance imaging (MRI) of the brain to exclude intracranial pathologies (brain tumor or previous stroke) and cognitive impairment. In the course of the study participants were excluded if they were not able to sleep long and deep enough during the first three nap sessions, i.e., stable NREM sleep (N2 or SWS) for at least 10 minutes (n = 14). Three participants were unable to undergo MRI, and were excluded from the study. In addition, one participant had to be excluded due to initially undisclosed use of a disqualifying medication, one participant due to impaired cognitive performance (e.g. VLMT recognition percentile rank score <5), one participant due to sleep apnea and one participant discontinued his participation, resulting in a sample of 33 patients who completed the experiment.

To ensure reliable results, 5 additional patients had to be excluded from analyses because they either exhibited only a very small number of SO events during analysis intervals following so-tDCS/sham, n=3) or their eigenfrequencies were assessed at 0.75 Hz, the same as the standardized frequency and were excluded on this basis (n=2). Therefore, 28 participants were included in the final analyses.

#### Baseline assessment

Comprehensive neuropsychological testing was conducted including memory (German version of Auditory Verbal Learning Test, AVLT [1]; Rey-Osterrieth Complex Figure Test [2]), working memory (digit span [3]), executive functions (Stroop color–word test [4]), verbal fluency (Regensburg Verbal Fluency Test [5]), processing speed and set shifting (Letter digit substitution test [4]) and verbal intelligence (Multiple choice vocabulary intelligence test, MWT-B [6]). The affective state at the time of the testing was assessed using the Positive and Negative Affect Schedule (PANAS [7]). Moreover, psychiatric comorbidity was monitored by Beck's Depression Inventory II [8]. See Table 1 for baseline characteristics.

Further, questionnaires regarding recent sleep habits (the German version of Morningness-Eveningness-Questionnaire [9]), sleep quality (Pittsburgh Sleep Quality Index [10]), daytime sleepiness (Epworth Sleepiness Scale [11]), and the Essen questionnaire on age and sleepiness (Essener Fragebogen Alter-und Schläfrigkeit [12]) were administered.

#### Sleep monitoring and scoring

During each nap session, an actiCAP EEG cap equipped with active recording electrodes placed according to the extended 10-20 international EEG system (FP1, FP2, AFz, F7, Fz, F8, FC5, FC1, FC2, FC6, C3, Cz, C4, T7, T8, CP5, CP1, CP2, CP6, P7, P3, Pz, P4, P8, O1, and O2). During the first two nap sessions (adaptation and eigenfrequency nap), F3 and F4 were additionally required, as these positions were used to determine individual SO frequency in each participant. All electrode recordings were referenced to an electrode attached to the nose and FCz electrode location was used as ground site. Data were collected at a sample rate of 500Hz and an online bandpass filter between 0.05 and 127 Hz. Impedances were kept below 10kΩ. Additionally, EMG at the chin and horizontal and vertical EOG were recorded according to standard sleep monitoring.

The sleep architecture, including time and proportion spent in different sleep stages, was determined based on polysomnographic criteria according to Rechtschaffen & Kales [13]. For this purpose, EEG data were down-sampled to 250 Hz and 30 sec epochs were scored visually in sleep stages 1, 2, 3, and 4 and REM sleep, epochs of wakefulness, or movement artifacts. Sleep stages 3 and 4 were summarized as slow wave sleep (SWS) in all following analyses. Stimulation epochs were not scored due to strong artifacts in the EEG signal. Likewise, corresponding epochs in the sham session were not scored to obtain comparable time and proportions of sleep stages (for each sham duration: 30 s, 2 min and 5 min).

To control for possible confounding effects, factors known to influence sleep quality (e.g. caffeine consumption, sleep duration, smoking) were assessed before each nap, and sleepiness and activation (Tiredness Symptoms Scale, TSS [14]; Visual Analog Scale, VAS [15]) in addition to the affective state (the Positive and Negative Affect Schedule, PANAS [7]) were likewise acquired before and after each nap. Participants were asked to avoid caffeinated drinks during at least the last 1.5 h before starting each nap session. To monitor habitual bedtimes and wake times three days prior the experimental nap sessions, sleep diaries and actigraphy (GT3X, ActiGraph, Pensacola, FL, USA) were used.

**So-tDCS**

The oscillatory current was applied by two battery-driven stimulators (DC-Stimulator; NeuroConn, Ilmenau, Germany) to bilateral frontal electrodes at sites F3 and F4 (8 mm in diameter and mounted into an actiCAP, Brain Products, Germany) with reference electrodes placed at each mastoid (ipsilateral; likewise 8mm in diameter). The stimulation was initiated by a trigger to ensure interhemispheric phase coherence and oscillated sinusoidally between zero and 262.5 µA, resulting in a maximum current density of 0.522 $mA/cm^2$ in each hemisphere. The electrode resistance was kept below 10 kΩ.

So-tDCS was started 4 minutes after the participant had entered stable NREM sleep stage 2 and was delivered in a blockwise manner either for 30 seconds, 2 minutes or 5 minutes, separated by stimulation-free interstimulus intervals of (at least) 90 seconds. These intervals were used to capture stimulation aftereffects, following rejection of the first 30 seconds to exclude the strong and long-lasting stimulation-induced drifts visible in our unfiltered online EEG signal from analysis (interval of analysis will be referred to hereafter as “1-min stimulation-free interval”). The number of stimulation trains (3–5 trains; 3 were required for inclusion in analysis) and the duration of the interstimulus intervals depended on the individual participant’s sleep, as sleep was monitored after each stimulation train and each train was only initiated during NREM sleep stage 2 or SWS. If the subject moved from sleep stage 2 (S2) or SWS to sleep stage 1 (S1), REM sleep, or wakefulness after a so-tDCS/sham train, the interstimulus interval was prolonged until the participant had re-entered S2 for at least 1 minute. This procedure was chosen to ensure state depend stimulation, since previous studies indicated that so-tDCS effects critically depend on ongoing brain state [16,17].

During the sham condition, stimulation electrodes were placed identical to the other conditions, but the tDCS device remained off. The same criteria as for the so-tDCS condition were applied for the sham condition: the first sham train was marked 4 minutes after onset of S2 and for subsequent sham trains, S2 or slow-wave sleep was required. In order to be able to compare the three different stimulation lengths with corresponding sham conditions, the one sham session was marked for each stimulation train length separately (30 s, 3 min and 5 min).

**Assessment of eigenfrequency**

To determine the individual peak SO frequency (termed eigenfrequency) during one nap session prior to experimental conditions, EEG data for S2 and SWS was epoched in overlapping (by 30sec) artifact-free segments of 60 sec. On each of these segments, a Hanning window was applied before calculating the power spectra (frequency resolution 0.001 Hz) using the Fast Fourier Transform (FFT). The following channels were selected for the analysis: Fp1, Fp2, AFz, Fz, F3 and F4. The FFT was averaged across all channels and smoothed using a moving average method (smooth percentage set at 0.06). Subsequently, mean power ($\mu V^2$) was calculated over the frequency range of 0.4 - 1.25 Hz and a Gaussian function was fit to the data to obtain the peak within the frequency range. To assess the eigenfrequency the FieldTrip toolbox [18] for MATLAB (The MathWorks) was used.

**Preprocessing**

All EEG preprocessing was carried out with MNE-Python v. 0.23 [19]. Data were filtered with a finite impulse response bandpass of 0.1-200 Hz. To remove line noise, notch filters were applied at 50 Hz and multiples thereof, up to the upper limit of the bandpass. Bad channels were algorithmically marked with the ANOAR package ANOAR package ([20], see Appendix), which identifies channels that do not correlate with neighboring channels for prolonged periods. An independent component analysis (ICA) was then performed with the Picard algorithm [21], decomposing the data into n components, where n is the number of good channels in a given recording. In order to remove ocular-related noise, ICA components which correlated highly with the EOG channels, as assessed by the iterated Z-score method in MNE-Python, were factored out of the data. SO detection was performed afterwards.

**Linear mixed effects models for Time-Frequency Representation (TFR)**

We used a Time-Frequency Representation (TFR) analysis in concert with linear mixed effects (LME) models to infer differences between stimulation conditions. After calculating TFR within the spindle band for each SO, baseline adjustment followed the z-score method, such that the mean and standard deviation of the time period -2.35 to -1.5s were used in a z-score transformation of the entire SO epoch. Correction for multiple comparisons was conducted with the permutation clustering approach with 1000 iterations. For each iteration, data were randomly shuffled, LME models were fit to the shuffled data, and the resulting test statistics were clustered with threshold-free clustering [22] (parameters: $h_0$=0, dh=0.2). The maximum and minimum points were selected from each iteration, and used to form the surrogate distribution of cluster weights. Clustering was then applied to the main LME test statistics, and the probability of cluster values under the null hypothesis were calculated from the surrogate distribution. We used an alpha of 0.05.

Despite our efforts to ensure interhemispheric phase coherence of the stimulation, a technical error with one of our stimulation devices occurred which caused the phase of the left and right stimulation sites to drift apart slowly at a rate of approximately .1 radian (5.7 degrees) per 30 seconds. This problem emerged in 12 of the 28 included participants. The effect this has on SO-spindle coupling is unknown, so we modeled it in the LME model as a random effect, specifically the interaction of a two-level Synchronicity variable (synchronous or not), a two-level Stimulation variable (sham or eigen-/standardized frequency stimulation), and a three level Duration variable (30s, 2min, 5min). We infer that any remaining, significant fixed effects are independent of whatever variance may have been caused by non-synchronous stimulation.

**ERPAC**

Comparisons between stimulation and sham conditions were conducted according to Voytek et al. [23]: Fisher's z-transform was applied to the rho values in order to normalise them. Then the sham condition's z-transform was subtracted from the stimulation conditions' z-transforms. These were then divided by their respective standard errors to derive z-scores of the difference between stimulation and sham condition ERPACs, allowing straightforward statistical inference. Multiple comparisons correction was carried out in the same fashion as the TFR analysis (see above).

**Phase of spindle maxima**

For the analysis of phase of spindle maxima, distributions of phases across the sham, eigen-, and standardized frequency conditions were summarized with their resultant vectors, which were calculated by taking the mean of the vector [cos(ϕ), sin(ϕ)] for each phase ϕ]. The direction of the resultant vector indicates the mean phase of a given condition, and the length of the vector is inversely related to the variance within the condition.

In addition to carrying out these procedures on the phases, we also did the same procedures on the participant mean phases, in order to assess more general stimulation effects on participants' SO-spindle coupling.

**Supplementary Figure**

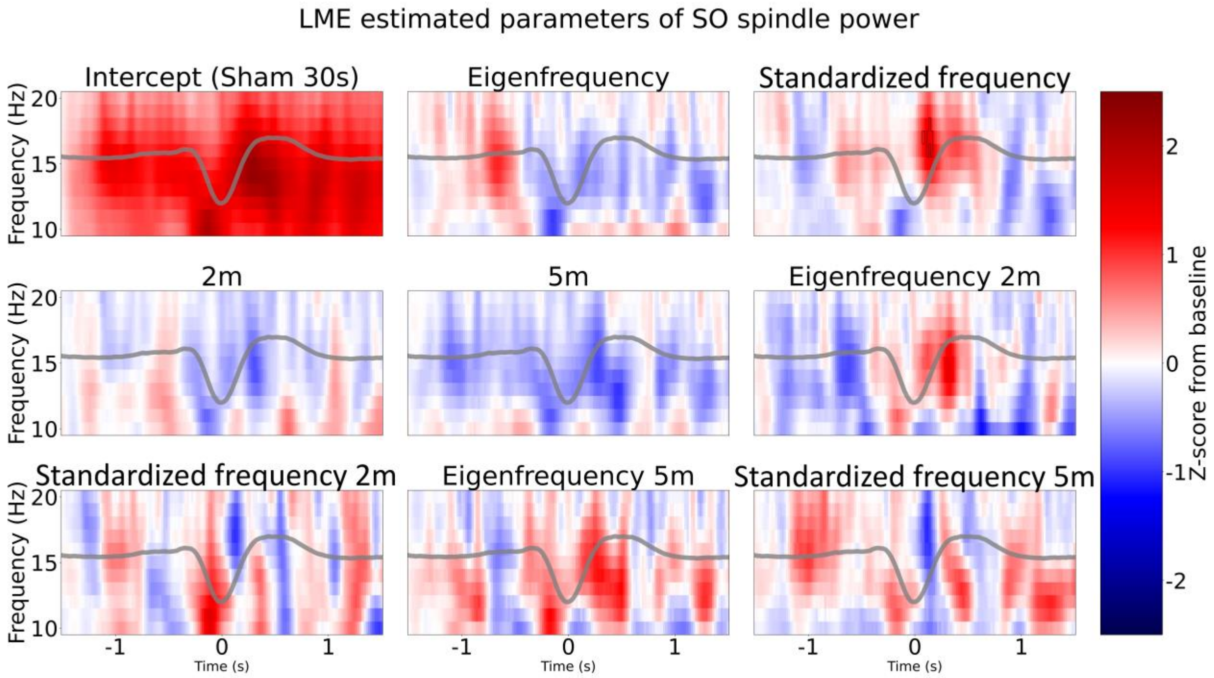


**Supplementary Figure 1: LME model parameter estimates for spindle band power during slow oscillations.** Each panel depicts the estimated parameters for a given fixed effect. The sham 30 s condition was chosen as the baseline, or intercept. Significance of parameter estimates, corrected for multiple comparisons at $p < 0.05$, are shown with the black lines. Averaged slow oscillations are superimposed in gray.

**Supplementary Table 1. Sleep architecture during each experimental condition.**

| 30 sec condition | | | | | |
|---|---|---|---|---|---|
| Sleep time (min) | sham stimulation | | eigenfrequency | | *p* |
| | Mean | SD | Mean | SD | |
| awake | 8.75 | 10.61 | 7.61 | 10.31 | 1.000 |
| NREM stage 1 | 24.98 | 15.06 | 28.88 | 18.97 | 1.000 |
| NREM stage 2 | 36.54 | 15.48 | 37.04 | 18.72 | 1.000 |
| NREM stage 3 | 2.63 | 4.68 | 1.73 | 4.25 | 1.000 |
| NREM stage 4 | 0.14 | 0.76 | 0.05 | 0.28 | 1.000 |
| REM | 1.16 | 3.41 | 0.96 | 3.12 | 1.000 |

| 30 sec condition | | | | | |
|---|---|---|---|---|---|
| Sleep time (min) | sham stimulation | | standardized frequency | | *p* |
| | Mean | SD | Mean | SD | |
| awake | 8.75 | 10.61 | 4.73 | 6.07 | 0.322 |
| NREM stage 1 | 24.98 | 15.06 | 28.07 | 18.96 | 1.000 |
| NREM stage 2 | 36.54 | 15.48 | 40.18 | 17.32 | 1.000 |
| NREM stage 3 | 2.63 | 4.68 | 3.37 | 7.57 | 1.000 |
| NREM stage 4 | 0.14 | 0.76 | 0.86 | 4.06 | 0.802 |
| REM | 1.16 | 3.41 | 1.57 | 3.71 | 1.000 |

| 30 sec condition | | | | | |
|---|---|---|---|---|---|
| Sleep time (min) | standardized frequency | | eigenfrequency | | *p* |
| | Mean | SD | Mean | SD | |
| awake | 4.73 | 6.07 | 7.61 | 10.31 | 0.742 |
| NREM stage 1 | 28.07 | 18.96 | 28.88 | 18.97 | 1.000 |
| NREM stage 2 | 40.18 | 17.32 | 37.04 | 18.72 | 1.000 |
| NREM stage 3 | 3.37 | 7.57 | 1.73 | 4.25 | 0.851 |
| NREM stage 4 | 0.86 | 4.06 | 0.05 | 0.28 | 0.637 |
| REM | 1.57 | 3.71 | 0.96 | 3.12 | 1.000 |

| 2 min condition | | | | | |
|---|---|---|---|---|---|
| Sleep time (min) | sham stimulation | | eigenfrequency | | *p* |
| | Mean | SD | Mean | SD | |
| awake | 8.29 | 10.41 | 5.81 | 7.59 | 0.736 |
| NREM stage 1 | 24.05 | 14.19 | 27.98 | 15.89 | 1.000 |
| NREM stage 2 | 30.73 | 15.2 | 35.15 | 15.15 | 0.917 |
| NREM stage 3 | 2.39 | 4.63 | 1.70 | 2.76 | 1.000 |
| NREM stage 4 | 0.14 | 0.76 | 0.70 | 3.56 | 0.970 |
| REM | 1.16 | 3.41 | 0.33 | 1.11 | 0.886 |

| 2 min condition | | | | | |
|---|---|---|---|---|---|
| Sleep time (min) | sham stimulation | | standardized frequency | | *p* |
| | Mean | SD | Mean | SD | |
| awake | 8.29 | 10.41 | 2.55 | 4.20 | 0.023* |
| NREM stage 1 | 24.05 | 14.19 | 26.39 | 17.02 | 1.000 |
| NREM stage 2 | 30.73 | 15.2 | 33.43 | 17.33 | 1.000 |
| NREM stage 3 | 2.39 | 4.63 | 3.71 | 5.00 | 0.748 |
| NREM stage 4 | 0.14 | 0.76 | 0.06 | 0.21 | 1.000 |
| REM | 1.16 | 3.41 | 1.02 | 3.51 | 1.000 |

| 2 min condition | | | | | |
|---|---|---|---|---|---|
| Sleep time (min) | standardized frequency | | eigenfrequency | | *p* |
| | Mean | SD | Mean | SD | |
| awake | 2.55 | 4.20 | 5.81 | 7.59 | 0.379 |
| NREM stage 1 | 26.39 | 17.02 | 27.98 | 15.89 | 1.000 |
| NREM stage 2 | 33.43 | 17.33 | 35.15 | 15.15 | 1.000 |
| NREM stage 3 | 3.71 | 5.00 | 1.70 | 2.76 | 0.252 |
| NREM stage 4 | 0.06 | 0.21 | 0.70 | 3.56 | 0.775 |
| REM | 1.02 | 3.51 | 0.33 | 1.11 | 1.000 |

| 5 min condition | | | | | |
|---|---|---|---|---|---|
| Sleep time (min) | sham stimulation | | eigenfrequency | | *p* |
| | Mean | SD | Mean | SD | |
| awake | 7.41 | 10.17 | 3.86 | 6.25 | 0.273 |
| NREM stage 1 | 21.95 | 13.20 | 21.19 | 12.99 | 1.000 |
| NREM stage 2 | 21.70 | 12.71 | 25.95 | 13.18 | 0.706 |
| NREM stage 3 | 1.63 | 3.93 | 3.59 | 8.44 | 0.567 |
| NREM stage 4 | 0.14 | 0.76 | 0.64 | 1.67 | 0.306 |
| REM | 0.86 | 2.35 | 0.64 | 2.16 | 1.000 |

| 5 min condition | | | | | |
|---|---|---|---|---|---|
| Sleep time (min) | sham stimulation | | standardized frequency | | *p* |
| | Mean | SD | Mean | SD | |
| awake | 7.41 | 10.17 | 3.91 | 6.17 | 0.296 |
| NREM stage 1 | 21.95 | 13.20 | 24.61 | 16.16 | 1.000 |
| NREM stage 2 | 21.70 | 12.71 | 26.63 | 14.01 | 0.519 |
| NREM stage 3 | 1.63 | 3.93 | 1.33 | 2.17 | 1.000 |
| NREM stage 4 | 0.14 | 0.76 | 0.15 | 0.68 | 1.000 |
| REM | 0.86 | 2.35 | 1.02 | 3.13 | 1.000 |

| 5 min condition | | | | | |
|---|---|---|---|---|---|
| Sleep time (min) | standardized frequency | | eigen frequency | | *p* |
| | Mean | SD | Mean | SD | |
| awake | 3.91 | 6.17 | 3.86 | 6.25 | 1.000 |
| NREM stage 1 | 24.61 | 16.16 | 21.19 | 12.99 | 1.000 |
| NREM stage 2 | 26.63 | 14.01 | 25.95 | 13.18 | 1.000 |
| NREM stage 3 | 1.33 | 2.17 | 3.59 | 8.44 | 0.407 |
| NREM stage 4 | 0.15 | 0.68 | 0.64 | 1.67 | 0.326 |
| REM | 1.02 | 3.13 | 0.64 | 2.16 | 1.000 |
